# Modeling effects of inter-group contact on links between population size and cultural complexity

**DOI:** 10.1101/2022.09.11.507470

**Authors:** Yotam Ben-Oren, Sarah Saxton Strassberg, Erella Hovers, Oren Kolodny, Nicole Creanza

## Abstract

Human populations rely on cultural artifacts and complex cumulative culture for their survival. Populations vary dramatically in the size of their tool repertoires, and the determinants of these cultural repertoire sizes have been the focus of extensive study in recent years. A prominent hypothesis, supported by computational models of cultural evolution, asserts that tool repertoire size increases with population size. However, not all empirical studies seeking to test this hypothesis have found such a correlation; this has led to a contentious and ongoing debate. As a possible resolution to this longstanding controversy, we suggest that accounting for even rare cultural migration events that allow sharing of knowledge between different-sized populations may help explain why a population’s size might not always predict its cultural repertoire size. Using an agent-based model to explore different assumptions about the effects of population size and migration on tool repertoires, we find that connectivity of one population to others, particularly to large populations, may significantly boost its tool repertoire size when population interactions lead to cultural exchange. Thus, two populations of identical size may have drastically different tool repertoire sizes, hinging upon their access to other groups’ knowledge. Intermittent contact between populations boosts cultural repertoire size and still allows for the development of unique tool repertoires that have limited overlap between populations.

## 1 Introduction

Human populations can accumulate massive repertoires of complex tools, artifacts, and technologies, varying in size and complexity[1, 2]. Many studies have attempted to explain this variation using computational models, laboratory experiments, and empirical analyses of hunter-gatherer and food-producing populations [3–12]. One prominent yet highly debated hypothesis predicts a positive correlation between population size and tool repertoire size, which is often used as a proxy of cultural complexity [13–17]. Different mechanisms can contribute to such an effect: for a given per capita innovation rate, large groups will invent more tools simply because they contain more innovators. Further, the presence of more learners can also reduce tool loss, facilitating cultural accumulation; smaller populations are at higher risk of losing technologies because fewer individuals know each tool[4, 18]. Additionally, contact between groups may allow for combination of cultural features (as explored in [8]), which may also increase the cultural repertoire. Since the 1960s [19], a number of findings have supported this hypothesis [5, 10, 20–23], some have provided conflicting evidence [9, 13, 24–26], and others suggest it might have more predictive power in certain contexts (i.e., in food-producing societies more than in hunter-gatherer groups) [27]. In this article, we use an agent-based model of cultural evolution to illustrate a simple scenario that could cause cultural repertoire sizes to deviate from the predictions of the population-size hypothesis even when the underlying assumptions hold: when there is inter-group contact, even when it occurs rarely, the cultural repertoire of a focal population could be influenced by those of the populations connected to it. Just as gene flow introduces new genetic variation and makes distant groups more genetically similar, connectivity allows sharing of novel tools between populations and promotes cultural overlap [28]. Previous researchers have noted that census population size might not accurately predict the cultural repertoire size; instead, they suggest that the size of the broader population that shares cultural information is more relevant than the census size of a sub-population, and populations that have moderate to high levels of contact should be considered part of the same population [17]. Here, we suggest that even rare events of cultural migration between otherwise distinct populations can have lasting effects on their cultural repertoires. Thus, investigating the contact networks of culturally distinct populations can play an important role in predicting cultural repertoire size. The effects of cultural exchange on tool repertoire size are related to both properties of individual populations and meta-population structure. For instance, the size of a neighboring population influences the focal population’s repertoire in two ways. First, larger populations have more tools to transmit. Second, for a given individual probability of migration, larger populations will produce more migrating individuals, so their tools might reach other groups more frequently [29]. We use an agent-based model to explore the combined effects of population size and migration on tool repertoire size. Specifically, we examine inter-group connectivity as a possible explanation both for variation in tool repertoire size between populations and for discrepancies among previous studies.

## 2 Methods

We used a simplified version of the agent-based model proposed in [8] (see https://github.com/CreanzaLab/ Cultural-Connectivity for more information on this article’s model and the code to replicate simulations). This model simulates accumulation and loss of tools over many generations; the rates of tool invention and tool loss are dependent on the number of individuals in a population. New tools are invented with a probability of *P*_*inv*_ per individual per time step. Each new tool is associated with a positive selection coefficient *s*, which is drawn from an exponential distribution with parameter *β*. We use the approximation that the probability of fixation of an adaptive allele is *s* [30] instead of explicitly simulating the establishment of the tool (i.e. its transmission within the population). Thus, we assume that a newly invented tool is established with probability *s*, and that it immediately reaches its equilibrium frequency in the population. Afterwards, we assume established tools can still be lost in the population with a probability *P*_*loss*_, which is divided by the number of individuals in the population. Thus, an isolated population will reach its equilibrium repertoire size when *P*_*inv*_ *β N* = *P*_*loss*_*/N*, where *N* is the population size. From this equation, for a given repertoire size of an interconnected population, we can calculate its “effective cultural population size”, that is, the population size at which an isolated population would be expected to have the same repertoire size. For simplicity, we did not consider in this model inter-dependence in invention, spread, or retention of tools whose functions are related. These are worthy of future exploration in the contexts of the proposed phenomena.

The parameter values we chose for this proof-of-principle model (*P*_*inv*_ = 0.001, *β* = 0.1, *P*_*loss*_ = 0.2) were chosen such that they would yield reasonable repertoire sizes for moderate population sizes. However, qualitatively similar results can be found for a wide parameter range.

Migration between populations occurs with a probability *P*_*mig*_ per individual. Migration events occur from the focal population to the neighbor and vice-versa and do not affect their population sizes. Each migrating individual carries with it a fraction *f* of its native population’s cultural repertoire. In the presented results, we set *f* to 0.5, however we found that changing *f* had virtually the same effect as changing *P*_*mig*_ (meaning the migration of two individuals carrying each 10% of their population’s tools had a very similar effect to the migration of one individual carrying 20% of the population’s tools). For each tool carried to another population, we assume the tool establishes in that population with the same probability *s* that was drawn from the exponential distribution described above when it was invented.

Repertoire sizes at equilibrium were calculated by averaging the repertoire sizes between time steps 50,000 (at which point repertoire size had already plateaued) and 100,000. Within the same time range, the average number of tools unique to the focal population was estimated by sampling the number of unique tools once every 100 time steps and averaging them.

## 3 Results

As proof of principle, we examined tool repertoire size of a simulated focal population of census size 200. The focal population was connected to a neighbor population of varying size (200 to 2000 individuals) with varying probabilities of migration (10^*−*6^ to 10^*−*5^ per individual, corresponding to between 0.0002 and 0.02 migration events per time step depending on the population size). For isolated populations, our model clearly predicts larger populations will have larger cultural repertoires. When migration is introduced, however, we find that the rate of migration has a pronounced effect on the repertoire size of the focal population of 200 individuals. When migration occurred between the focal and the neighbor populations, larger population size in the neighbor led to an increased cultural repertoire in the focal population. This concept can be described in terms of an ‘effective cultural population size’: the size of an isolated population that would be expected to have this cultural repertoire size at equilibrium (Figure 1A). Similarly, while even low connectivity led to an increase in effective cultural population size, this effect was stronger as the connectivity increased. These results demonstrate that a group of fixed size can have a tool repertoire characteristic of a much larger group, depending on the composition of its contact network.

**Figure 1:**
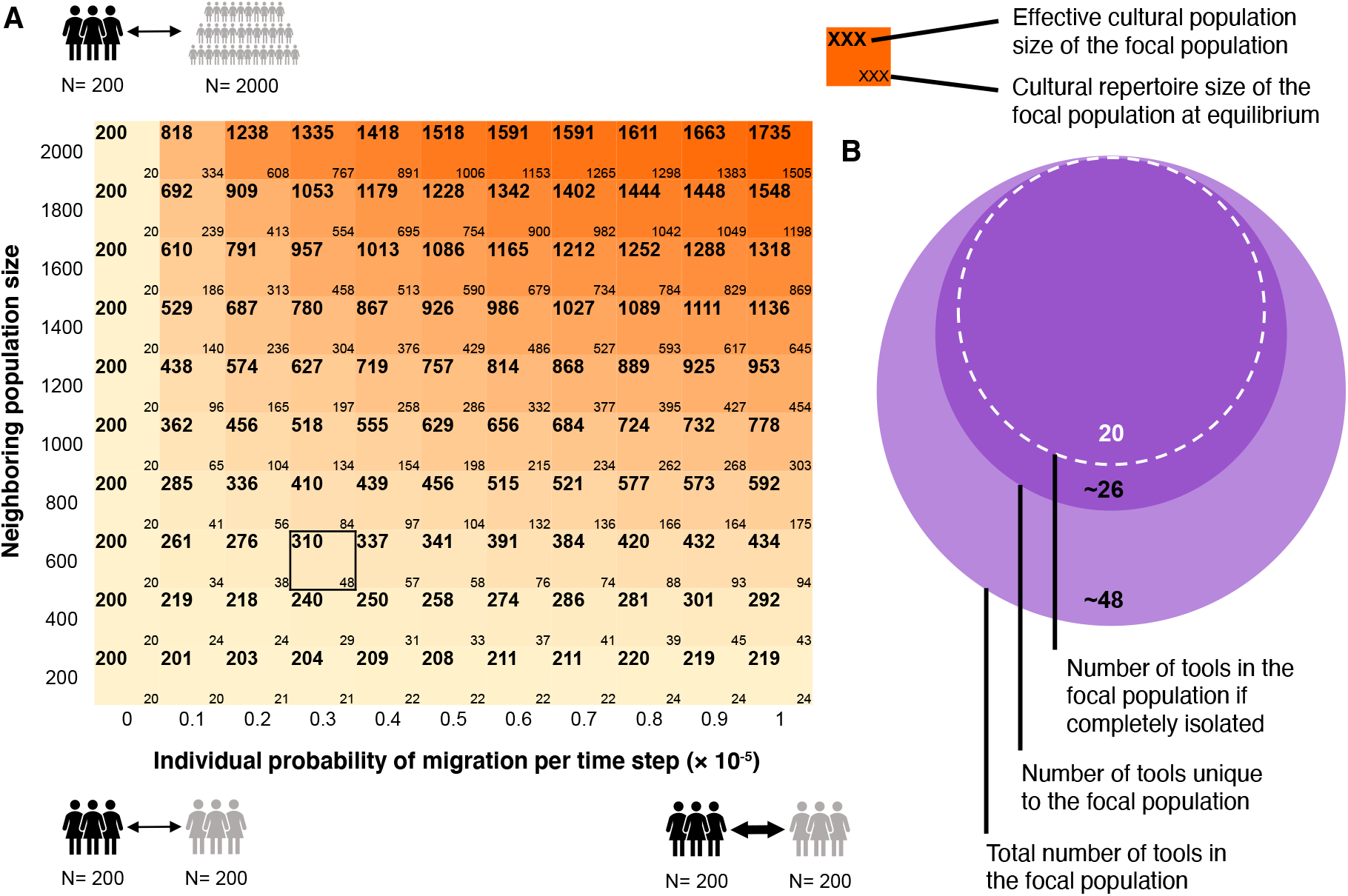
Inter-group contact may increase cultural repertoire size. In panel *A*, the heatmap indicates the cultural repertoire of a population of size 200 (illustrated by the black figures) that has relatively rare contact with one other population. In this population, we show that both migration rate (varying on the *x*-axis between 10^*−*6^ and 10^*−*5^ per individual) and the size of the neighboring population (varying on the *y*-axis between 200 and 2000, illustrated by the grey figures) can contribute to the population’s repertoire size at equilibrium (bottom numbers in cells) and thus to its effective cultural population size (top numbers). The effective cultural population size is the number of individuals in an isolated population that is expected to have a tool repertoire of the same size as the connected population of 200 individuals. Higher migration rates increase the effective population size, but larger neighboring populations have an even more dramatic effect. In panel *B*, we illustrate that connectivity to other populations not only increases cultural repertoire, but can also increase the number of tools that are unique to the focal population (in this case, a population of 200 individuals, connected to a population of 600 individuals with an individual probability of migration of 0.3 · 10^*−*5^)

This phenomenon accounts for an additional observation: we found that, depending on the specific loss mechanism used, when populations are highly connected (beyond the migration values shown in the figure) their effective cultural population size can exceed the sum of their population sizes.

Furthermore, in Figure 1B we show that, in many cases, connectivity to other populations not only increases the focal population’s overall repertoire, but surprisingly can also increase the number of tools that are unique to it, i.e. those tools found only in one population and not the other at a given time. In this example (corresponding to the cell framed in black in Figure 1A), when a population of size 200 is connected to a neighboring population of size 600, with an individual rate of migration of 3 · 10^*−*6^, its average repertoire size increases from 20 tools to ∼ 48, of which ∼ 26 are unique to it.

In Figure 1, migrating individuals each carry a fraction of the tool repertoire of their population. Thus, individuals leaving larger populations carry larger tool repertoires. We also tested the effects of a different assumption, that individuals migrate carrying with them a constant number of tools regardless of the repertoire size of their population. With this assumption, we observe qualitatively similar results to those in the heatmap in Figure 1A: the cultural repertoire of a population of size 200 increases both when it is connected to a larger population and when those connections occur more often. (The code and figures for all simulations are available at https://github.com/CreanzaLab/Cultural-Connectivity).

## 4 Discussion

Our results indicate that a population’s tool repertoire size increases with access to other populations’ cultural repertoires. In our model, each individual has the same probability to migrate, which means that a focal population has more encounters with individuals from larger populations. Furthermore, each migrating individual carries a given fraction of its population’s repertoire, meaning that those from larger populations carry more tools. Thus, populations that communicate with individuals from large groups experience an influx of novel tools, even when these interactions occur rarely.

When a small population is in contact with a larger population, one might expect that, culturally, it would simply become a “satellite population” of the larger group, carrying primarily some subset of the large population’s repertoire. Interestingly, this is not always the case. Contact with a larger population can not only increase a population’s absolute repertoire size, but also increase the number of tools that are unique to the focal population. This is because, first, some of the tools that are transmitted to the focal population are subsequently lost in the neighbor and thus become unique tools in the focal population. Second, the relative population sizes may lead to asymmetric migration rates, which keeps the smaller population distinct: migration rate is defined per individual in our model, and there are thus few individuals leaving the small population. Accordingly, most of the tools that originate in the small population remain unique to it.

Interestingly, we find that for very high migration rates the effective cultural population sizes of two interconnected populations can be larger than the sum of their population sizes. This result, while not intuitive, can be understood by considering the mechanisms of migration and tool loss. When migration rate is very high, most tools are shared between the populations. Thus, whenever a tool is lost in one population, it is reintroduced rapidly by migration from the other. For a tool to be lost from the meta-population it thus needs to be lost in both populations nearly simultaneously. In our model, the probability of tool loss in a population sized 2*N* is *P*_*loss*_*/*2*N*, which is larger than the probability of simultaneous loss in two *N* sized populations, which is (*P*_*loss*_*/N*)^2^. Under these assumptions, as populations are more interconnected, it becomes very rare for tools to be lost in the meta-population. We call this phenomenon “cultural rescue”, and believe it may play a meaningful role in the cultural dynamics of real meta-populations. Importantly, however, this phenomenon depends on the mechanisms by which tools are lost. In the version of the model used here, the “cultural rescue effect” appears because we implicitly assume that the probability that two individuals from the same population lose a given tool is higher than the probability that two individuals from different populations lose that tool simultaneously. In other words, individual-level loss events of a tool within the same population are more interdependent than loss events across populations. An alternative model in which tools are lost from a population when they are independently lost by all individuals would lead the relationship between population size and the probability of loss to be exponentially decreasing. In such a model, very high migration rates between the populations would drive their effective cultural population sizes to simply approach the sum of their population sizes, and the cultural rescue effect would not appear. A further complication may arise if cultural loss is not considered a completely stochastic phenomenon, but rather is more likely, for example, when environmental conditions make certain tools less useful [8]. In such a case, it is reasonable that highly connected populations would also share other factors—such as environmental conditions—that would couple their loss dynamics, limiting the impact of the cultural rescue effect.

We suggest that empirical studies to examine overlap between populations’ repertoires and reconstruct common origins of their tools and practices may help elucidate underlying evolutionary mechanisms of tool repertoire size variation. However, detecting infrequent contact events between populations may be extremely difficult. Here we demonstrate that, especially when a population of interest interacts with a large population, even rare migration events can contribute to its cultural complexity, obscuring the predicted links between its population size and the size of its cultural repertoire. Moreover, although one might expect that population connectivity would lead to highly homogenized cultural repertoires [4, 31, 32], here we demonstrate that it is possible that the two interconnected populations would have increased cultural repertoires which are distinct from one another.

## Acknowledgements

NC and SSS received funding from Vanderbilt University and the VU Summer Research Program. OK and YBO are supported by the US–Israel Binational Science Foundation (BSF) and by the Israeli Science Foundation (ISF; 1826/20). We thank Joe Henrich, Liran Carmel, and members of the Creanza and Kolodny labs for feedback.

## Notes

https://github.com/CreanzaLab/Cultural-Connectivity

### Competing Interest Statement

The authors have declared no competing interest.

## References

[1] R. Boyd and P. Richerson. Culture and the evolution of human cooperation. Philosophical Transactions of the Royal Society B: Biological Sciences, 364:3281–3288, 2009.

[2] M. Tomasello. The human adaptation for culture. Annual Review of Anthropology, 28:509–529, 1999.

[3] Briggs Buchanan, Michael O’Brien, and Mark Collard. Drivers of technological richness in prehistoric texas. Archaeological and Anthropological Sciences, 8(3):625–634, 2016.

[4] Joseph Henrich. Demography and cultural evolution: how adaptive cultural processes can produce maladaptive losses: the tasmanian case. American antiquity, 69:197–214, 2004.

[5] M. Kline and R. Boyd. Population size predicts technological complexity in oceania. Proceedings of the Royal Society B, 277:2559–2564, 2010.

[6] M. Collard, B. Buchanan, and M. O’Brien. Population size as an explanation for patterns in the paleolithic archaeological record. Current Anthropology, 54:S388–S396, 2013.

[7] M. Derex, M. Beugin, B. Godelle, and M. Raymond. Experimental evidence for the influence of group size on cultural complexity. Nature, 503:389–391, 2013.

[8] O. Kolodny, N. Creanza, and M. Feldman. Evolution in leaps: The punctuated accumulation and loss of cultural innovations. Proceedings of the National Academy of Sciences, 112:E6762–9, 2015.

[9] C. Andersson and D. Read. Group size and cultural complexity. Nature, 511:E1, 2014.

[10] R. Bentley and M. O’Brien. The selectivity of social learning and the tempo of cultural evolution. Journal of Evolutionary Psychology, 9:125–141, 2011.

[11] S. Shennan. Demography and cultural innovation: a model and its implications for the emergence of modern human culture. Cambridge Archaeological Journal, 11:5–16, 2001.

[12] G. Carlino, S. Chatterjee, and R. Hunt. Urban density and the rate of invention. Journal of Urban Economics, 61:389–419, 2007.

[13] K. Vaesen, M. Collard, R. Cosgrove, and W. Roebroeks. Population size does not explain past changes in cultural complexity. Proceedings of the National Academy of Sciences, 113:E2241–E2247, 2016.

[14] N. Fay, N. De Kleine, B. Walker, and C. Caldwell. Increasing population size can inhibit cumulative cultural evolution. Proceedings of the National Academy of Sciences, 116:6726–6731, 2019.

[15] J. Martens. Scenarios where increased population size can enhance cumulative cultural evolution are likely common. Proceedings of the National Academy of Sciences, 116:17160, 2019.

[16] K. Vaesen, M. Collard, R. Cosgrove, and W. Roebroeks. Reply to henrich et al.: The tasmanian effect and other red herrings. Proceedings of the National Academy of Sciences, 113:E6726–E6727, 2016.

[17] J. Henrich and et al. Understanding cumulative cultural evolution. Proceedings of the National Academy of Sciences, 113:E6724–E6725, 2016.

[18] Yotam Ben-Oren, Oren Kolodny, and Nicole Creanza. Cultural specialization as a double-edged sword. bioRxiv, page doi:10.1101/2022.04.22.489202, 2022.

[19] R. Carneiro. On the relationship between size of population and complexity of social organization. Southwestern Journal of Anthropology, 23:234–243, 1967.

[20] L. Premo and S. Kuhn. Modeling effects of local extinctions on culture change and diversity in the paleolithic. PLoS One, 5:e15582, 2010.

[21] M. Kempe and A. Mesoudi. An experimental demonstration of the effect of group size on cultural accumulation. Evolution and Human Behavior, 35:285–290, 2014.

[22] M. Muthukrishna, B. Shulman, V. Vasilescu, and J. Henrich. Sociality influences cultural complexity. Proceedings of the Royal Society B, 281:20132511, 2014.

[23] L. Bettencourt, J. Lobo, and D. Strumsky. Invention in the city: Increasing returns to patenting as a scaling function of metropolitan size. Research Policy, 36:107–120, 2007.

[24] M. Collard, A. Ruttle, B. Buchanan, and M. O’Brien. Population size and cultural evolution in nonindustrial food-producing societies. PLoS One, 8:e72628, 2013.

[25] M. Nelson and et al. Resisting diversity: a long-term archaeological study. Ecology and Society, 16, 2011.

[26] N. Fay, N. De Kleine, B. Walker, and C. Caldwell. Population size and cumulative cultural evolution: fewer heads can be better than many. 2019.

[27] L. Fogarty and N. Creanza. The niche construction of cultural complexity: interactions between innovations, population size and the environment. Phil. Trans. Roy. Soc. B, 372, 2017.

[28] N. Creanza, O. Kolodny, and M. Feldman. Greater than the sum of its parts? modelling population contact and interaction of cultural repertoires. Journal of the Royal Society Interface, 14, 2017.

[29] R. Boyd and P. Richerson. Voting with your feet: Payoff biased migration and the evolution of group beneficial behavior. Journal of Theoretical Biology, 257:331–339, 2009.

[30] Motoo Kimura. On the probability of fixation of mutant genes in a population. Genetics, 47(6): 713, 1962.

[31] A. Migliano and et al. Hunter-gatherer multilevel sociality accelerates cumulative cultural evolution. Science Advances, 6:eaax5913, 2020.

[32] M. Derex and R. Boyd. Partial connectivity increases cultural accumulation within groups. Proceedings of the National Academy of Sciences, 113:2982–2987, 2016.

